## Supplemental Figures and Tables for "Niche partitioning facilitates coexistence of closely related gut bacteria"

### Supplemental Tables:

**Supplementary Table 1:** List of primers used in this study: the sample-specific barcodes used in the primers for the second PCR of the amplicon sequencing are highlighted in grey.

#### Amplicon Sequencing PCR I:

| |
|---|
| **F**: CGTACGTAGACGGCCAGTATGCCNGAAATGCCRGARGTTGA |
| **R**: GACTGACTGCCTATGACGACTAARCGATAYTTRCCYTCCATRCG |

#### Amplicon Sequencing PCR II:

| | |
|---|---|
| **B1F/B1R** | CAAGCAGAAGACGGCATACGAGATTAAGAGGCGTCTCGTGGGCTCGGAGATGTGTATAAGAGACAGATACGTACGTA<br>GACGGCCAGT/AATGATACGGCGACCACCGAGATCTACACGCCTCTTATCGTCGGCAGCGTCAGATGTGTATAAGAG<br>ACAGGACTGACTGCCTATGACG |
| **B2F/B2R** | CAAGCAGAAGACGGCATACGAGATGCGAATTCGTCTCGTGGGCTCGGAGATGTGTATAAGAGACAGTACGTACGTAG<br>ACGGCCAGT/AATGATACGGCGACCACCGAGATCTACACCAGCGTATTCGTCGGCAGCGTCAGATGTGTATAAGAGA<br>CAGTGACTGACTGCCTATGACG |
| **B3F/B3R** | CAAGCAGAAGACGGCATACGAGATACTGAGCTGTCTCGTGGGCTCGGAGATGTGTATAAGAGACAGGCGTACGTAGA<br>CGGCCAGT/AATGATACGGCGACCACCGAGATCTACACGAATTCGCTCGTCGGCAGCGTCAGATGTGTATAAGAGAC<br>AGGTGACTGACTGCCTATGACG |
| **B4F/B4R** | CAAGCAGAAGACGGCATACGAGATTTAGGCACGTCTCGTGGGCTCGGAGATGTGTATAAGAGACAGCGTACGTAGAC<br>GGCCAGT/AATGATACGGCGACCACCGAGATCTACACAGCTCAGTTCGTCGGCAGCGTCAGATGTGTATAAGAGACA<br>GAGTGACTGACTGCCTATGACG |
| **B5F/B5R** | CAAGCAGAAGACGGCATACGAGATCTCCGATTGTCTCGTGGGCTCGGAGATGTGTATAAGAGACAGATACGTACGTA<br>GACGGCCAGT/AATGATACGGCGACCACCGAGATCTACACGTGCCTAATCGTCGGCAGCGTCAGATGTGTATAAGAG<br>ACAGGACGACTGCCTATGACG |
| **B6F/B6R** | CAAGCAGAAGACGGCATACGAGATTTCACAGGGTCTCGTGGGCTCGGAGATGTGTATAAGAGACAGTACGTACGTAG<br>ACGGCCAGT/AATGATACGGCGACCACCGAGATCTACACACGTAAGGTCGTCGGCAGCGTCAGATGTGTATAAGAGA<br>CAGTGACTGACTGCCTATGACG |
| **B7F/B7R** | CAAGCAGAAGACGGCATACGAGATCAGAGGTAGTCTCGTGGGCTCGGAGATGTGTATAAGAGACAGGCGTACGTAGA<br>CGGCCAGT/AATGATACGGCGACCACCGAGATCTACACCCTGTGAATCGTCGGCAGCGTCAGATGTGTATAAGAGAC<br>AGGTGACTGACTGCCTATGACG |
| **B8F/B8R** | CAAGCAGAAGACGGCATACGAGATGGCATCATGTCTCGTGGGCTCGGAGATGTGTATAAGAGACAGCGTACGTAGAC<br>GGCCAGT/AATGATACGGCGACCACCGAGATCTACACTACCGAGTTCGTCGGCAGCGTCAGATGTGTATAAGAGACA<br>GAGTGACTGACTGCCTATGACG |
| **B9F/B9R** | CAAGCAGAAGACGGCATACGAGATTATGACCGGTCTCGTGGGCTCGGAGATGTGTATAAGAGACAGATACGTACGTA<br>GACGGCCAGT/AATGATACGGCGACCACCGAGATCTACACGGAATGCATCGTCGGCAGCGTCAGATGTGTATAAGAG<br>ACAGGACTGACTGCCTATGACG |
| **B10F/B10R** | CAAGCAGAAGACGGCATACGAGATATACGCTGGTCTCGTGGGCTCGGAGATGTGTATAAGAGACAGTACGTACGTAG<br>ACGGCCAGT/AATGATACGGCGACCACCGAGATCTACACAATCGGAGTCGTCGGCAGCGTCAGATGTGTATAAGAGA<br>CAGTGACTGACTGCCTATGACG |
| **B11F/B11R** | CAAGCAGAAGACGGCATACGAGATACGTTCTCGTCTCGTGGGCTCGGAGATGTGTATAAGAGACAGGCGTACGTAGA<br>CGGCCAGT/AATGATACGGCGACCACCGAGATCTACACGAGAACGTTCGTCGGCAGCGTCAGATGTGTATAAGAGAC<br>AGGTGACTGACTGCCTATGACG |
| **B12F/B12R** | CAAGCAGAAGACGGCATACGAGATAATTGGCCGTCTCGTGGGCTCGGAGATGTGTATAAGAGACAGCGTACGTAGAC<br>GGCCAGT/AATGATACGGCGACCACCGAGATCTACACGGCCAATT<br>TCGTCGGCAGCGTCAGATGTGTATAAGAGACAG AGTGACTGACTGCCTATGACG |
| **B13F/B13R** | CAAGCAGAAGACGGCATACGAGATCATGGCATGTCTCGTGGGCTCGGAGATGTGTATAAGAGACAGATACGTACGTA<br>GACGGCCAGT/AATGATACGGCGACCACCGAGATCTACACTTCGCATCTCGTCGGCAGCGTCAGATGTGTATAAGAG<br>ACAGGACTGACTGCCTATGACG |
| **B14F/B14R** | CAAGCAGAAGACGGCATACGAGATACTCGGTAGTCTCGTGGGCTCGGAGATGTGTATAAGAGACAGTACGTACGTAG<br>ACGGCCAGT/AATGATACGGCGACCACCGAGATCTACACATGCCATGTCGTCGGCAGCGTCAGATGTGTATAAGAGA<br>CAGTGACTGACTGCCTATGACG |
| **B15F/B15R** | CAAGCAGAAGACGGCATACGAGATGAGTTCCAGTCTCGTGGGCTCGGAGATGTGTATAAGAGACAGGCGTACGTAGA<br>CGGCCAGT/AATGATACGGCGACCACCGAGATCTACACTGATCACGTCGTCGGCAGCGTCAGATGTGTATAAGAGAC<br>AGGTGACTGACTGCCTATGACG |

#### qPCR:

| | |
|---|---|
| **F:** GCAACCTGCCCTWTAGCTTG | Ref: Kešnerová et al. (2017). |
| **R:** GCCCATCCTKTAGTGACAGC | Ref: Kešnerová et al. (2017). |

**Supplementary Table 2:** COG functional categories.

| COG Code | COG Function |
|---|---|
| J | Translation, ribosomal structure and biogenesis |
| A | RNA processing and modification |
| K | Transcription |
| L | Replication, recombination and repair |
| B | Chromatin structure and dynamics |
| D | Cell cycle control, cell division, chromosome partitioning |
| Y | Nuclear structure |
| V | Defense mechanisms |
| T | Signal transduction mechanisms |
| M | Cell wall/membrane/envelope biogenesis |
| N | Cell motility |
| Z | Cytoskeleton |
| W | Extracellular structures |
| U | Intracellular trafficking, secretion, and vesicular transport |
| O | Posttranslational modification, protein turnover, chaperones |
| X | Mobilome: prophages, transposons |
| C | Energy production and conversion |
| G | Carbohydrate transport and metabolism |
| E | Amino acid transport and metabolism |
| F | Nucleotide transport and metabolism |
| H | Coenzyme transport and metabolism |
| I | Lipid transport and metabolism |
| P | Inorganic ion transport and metabolism |
| Q | Secondary metabolites biosynthesis, transport and catabolism |
| R | General function prediction only |
| S | Function unknown |

### Supplemental Figures:

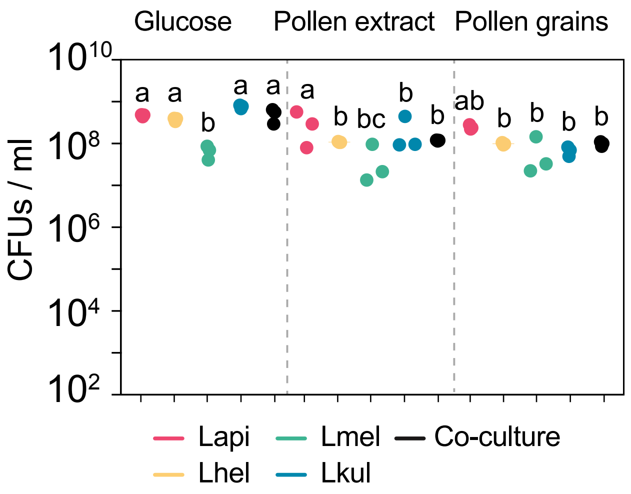


**Supplementary Figure 1.** Colony forming units (CFUs) per ml of culture after 24 h of growth of the four species in mono-cultures (n=3) or in co-culture (n=3) in the presence of 2% (w/v) glucose (G), 10% pollen extract (PE), or 10% pollen grains (PG). Statistical differences: ANOVA with Tuckey post-hoc test (BH correction), represented by different letters.

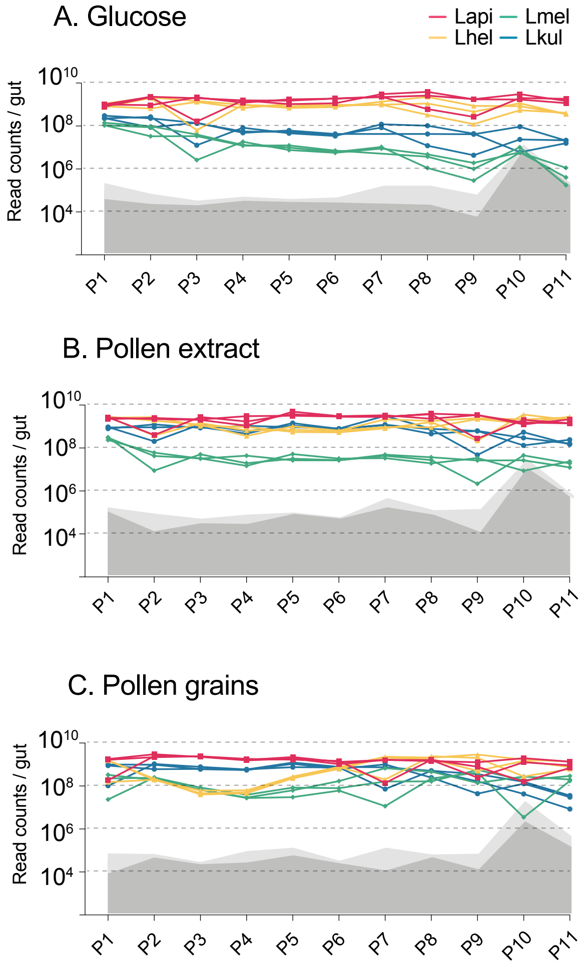


**Supplementary Figure 2: Second *in vitro* transfer experiment**. Changes in total bacterial abundance of the four species across the 11 serially passaged co-cultures in minimal medium supplemented with either 2% (w/v) glucose **(A),** 10% pollen extract (**B**), or 10% pollen grains **(C).** The absolute abundance of each species was determined by multiplying the total number of CFUs with the proportion of each strain in a given sample as based on amplicon sequencing. Grey areas (light grey = 95% CI) represent the limit of detection as explained in Figure 1 (see method).

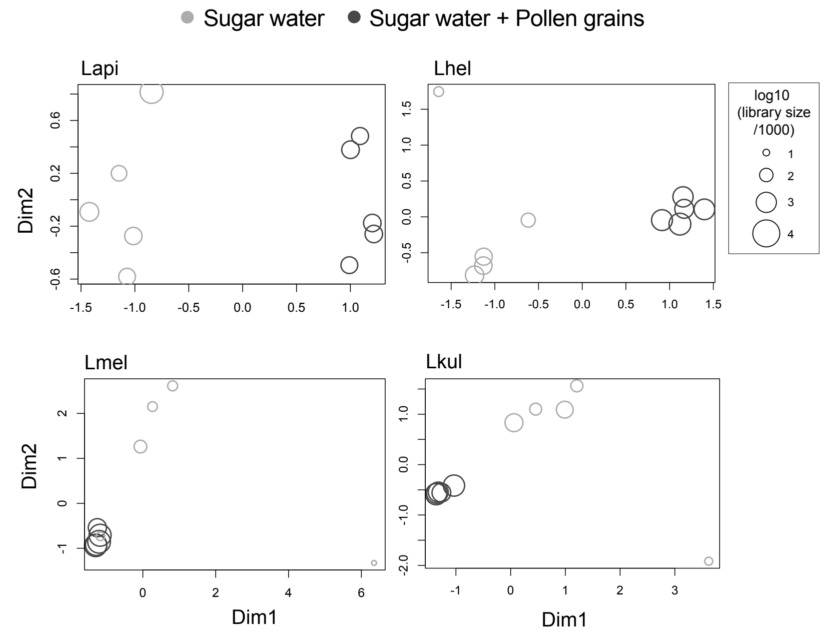


**Supplementary Figure 3. MDS plots of *in vivo* RNA-seq samples.** Counts per million (cpm) were calculated for each sample (n=5) and visualized using multidimensional scaling (MDS) plots. X- and y-axis axes show first and second MDS dimension, respectively. Shapes size correspond to the samples libraries size. A few samples of the SW treatment did not cluster with the other replicates (two samples for Lmel, and one sample for each Lhel and Lkul), in part because relatively few reads mapped to the reference genomes of these strains.

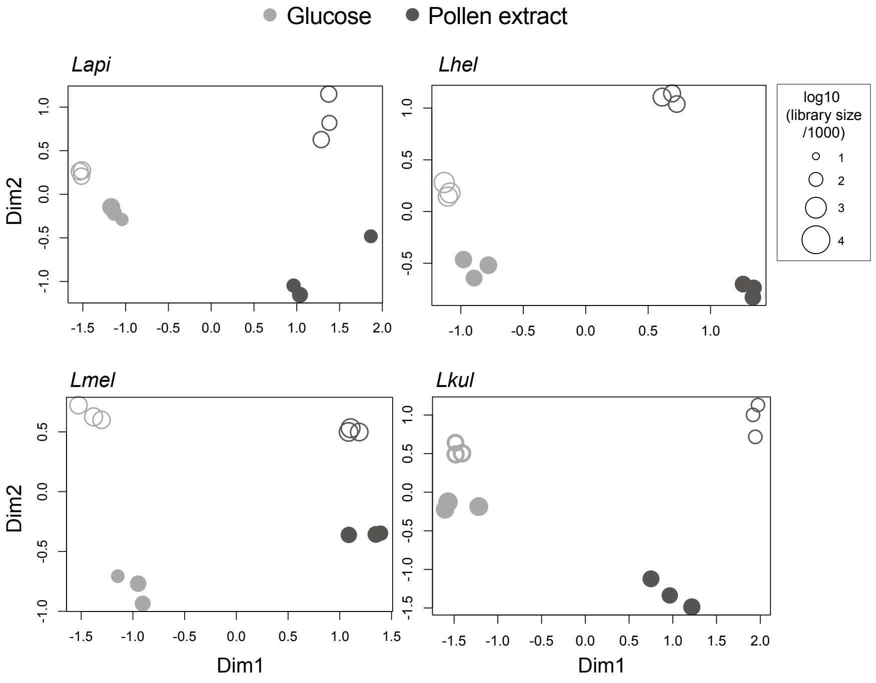


**Supplementary Figure 4**. **MDS plots of *in vitro* RNA-seq samples.** Counts per million (cpm) were calculated for each sample (n=3) and visualized using multidimensional scaling (MDS) plots. X- and y-axis axes show first and second MDS dimension, respectively. Filled shapes represent mono-culture samples and empty shapes represent co-culture samples. Shapes size corresponds to the libraries size of that sample.

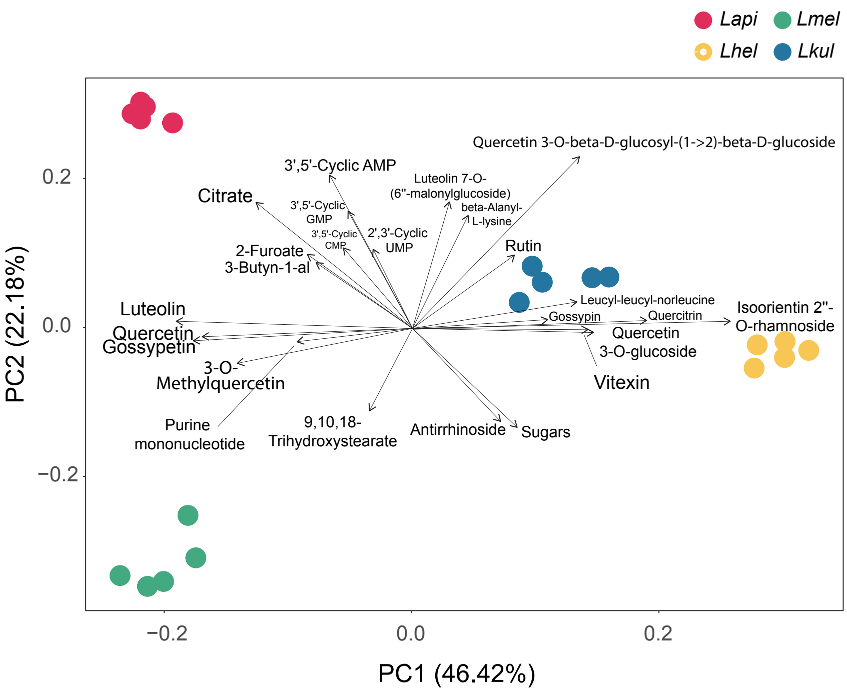


**Supplementary Figure 5**: **PCA *in vitro* metabolomics**. Principal component analysis (PCA) of the metabolome profile of each species based on the $log_2FC$ values calculated between the two time-points for each ion. The larger the distance between species on the PCA axes, the more they differ in their metabolome profiles. The arrows, i.e. the environmental vectors, point in the direction of the maximum correlation with the environmental variable, i.e. the ions. The ions on the tip of the longest arrows are the ones that explain the most of the distribution of the data within the PCA. Only the top 24 ions explaining the data distribution are displayed.

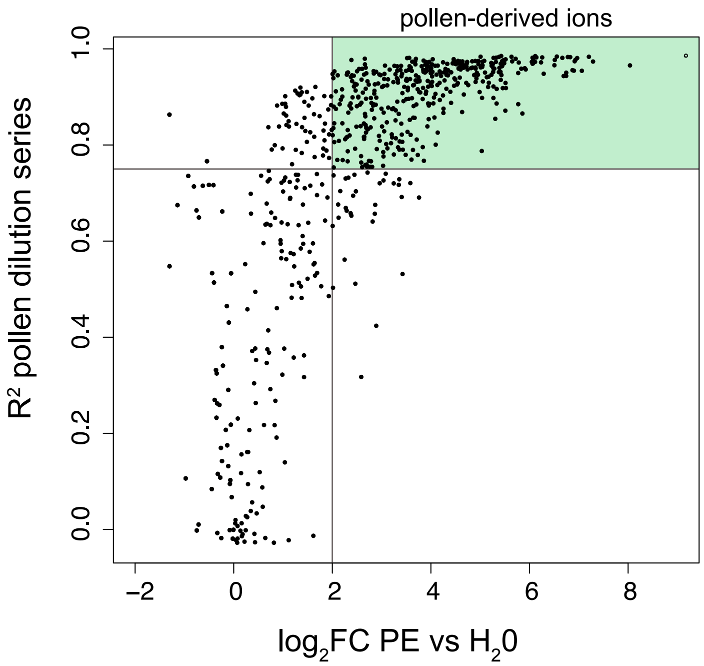


**Supplementary Figure 6**: **Definition of pollen-derived ions**. Volcano plot displaying $R^2$ values obtained from the pollen dilution series regression lines and the $log_2FC$ calculated between undiluted pollen extract and water. The lines represent the thresholds that we set to define an ion as pollen-derived: log2FC > 2 and $R^2 > 0.75$. Within the light green area are included the ions that we consider pollen-derived (n = 406).

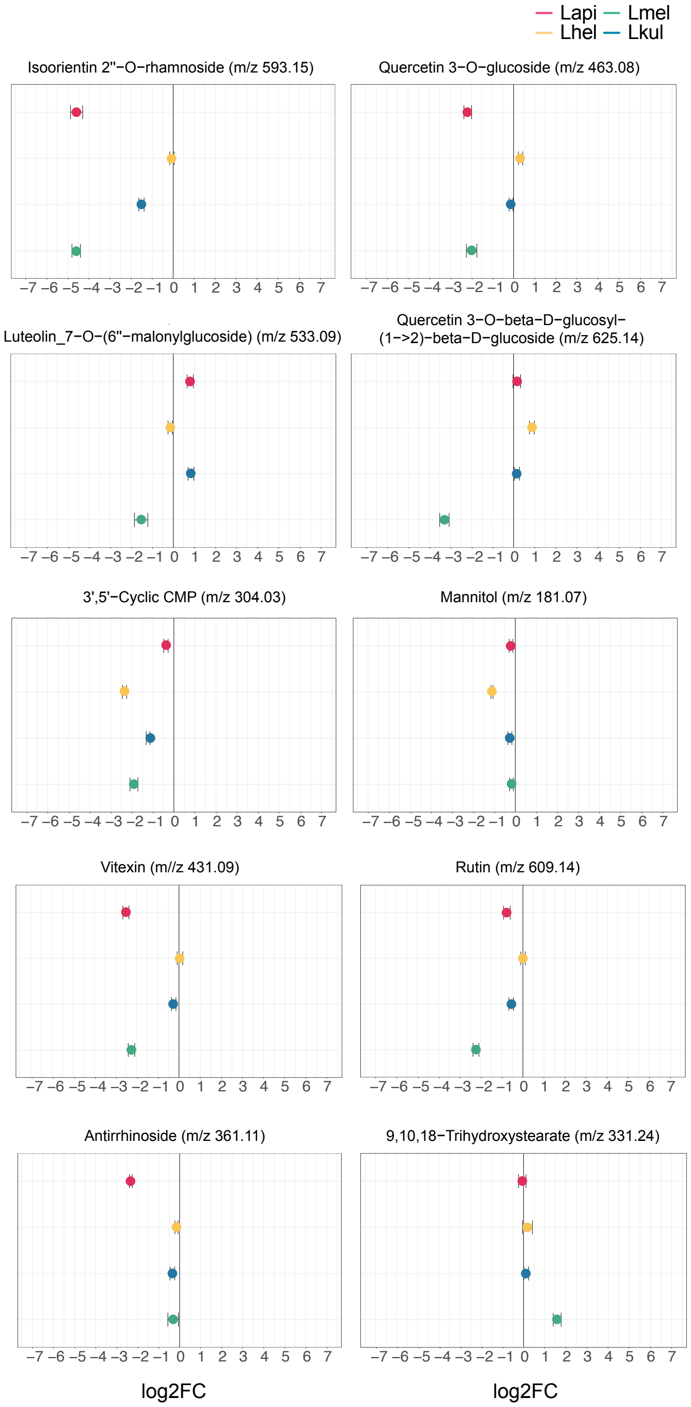


**Supplementary Figure 7**: **Untargeted metabolomics: key metabolites discussed in the main text**. *In vitro* metabolomics of spent medium of the four species grown in cfMRS + PE for 16 hours. The $log_2FC$ was obtained comparing the ion intensities at the end and at the beginning of the experiment.

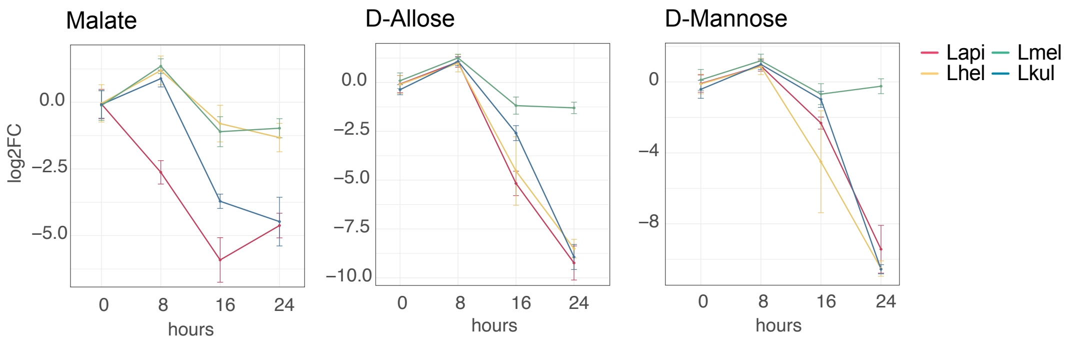


**Supplementary Figure 8: GC-MS detection of key metabolites over time**. $Log_2FC$ relative to T0 is plotted. Time is reported in hours. For m/z values see Supplementary Dataset 8.

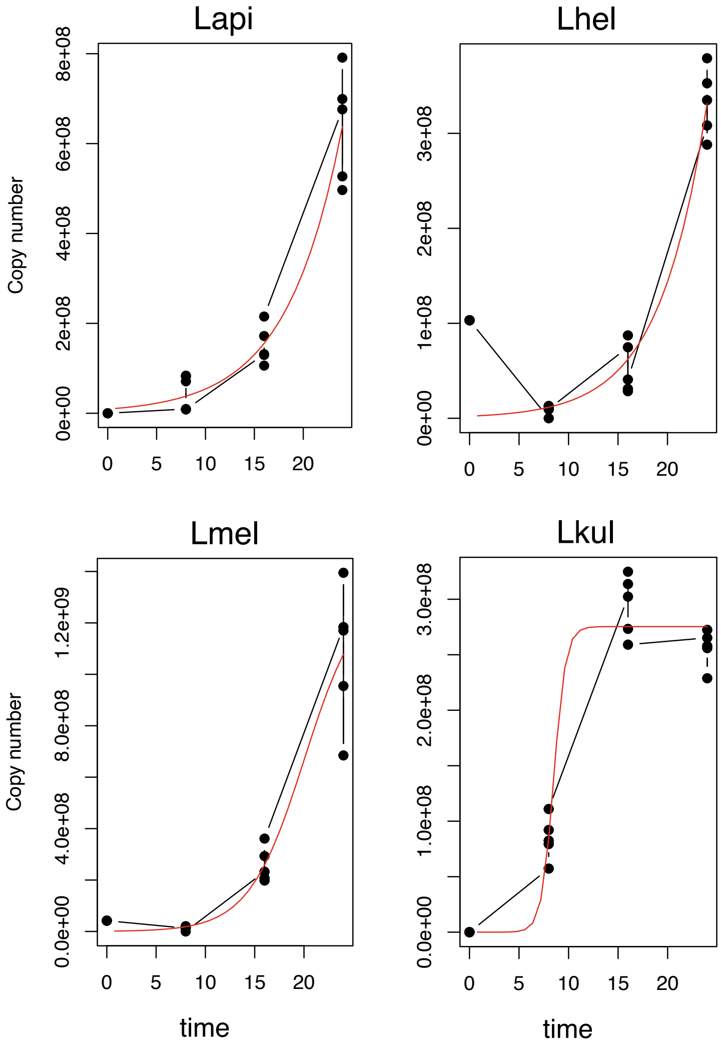


**Supplementary Figure 9: Logistic regression growth curve of the four species.** Growth-curve data were obtained for the four species at the four time-points included in the second metabolomics experiment (growth in presence of pollen extract) by qPCR (copy number) and fitted to a standard form of the logistic equation. Each point represents one replicate (n=5).
